# CMS: Achieving Uniform and High-Quality Sequencing across Challenging Non-canonical Genomic Regions

**DOI:** 10.64898/2026.04.24.720553

**Authors:** Qigang Li, Lei Liu, Qunting Lin, Xu Dan, Yuzhou Jiang, Yanli Wei, ManMan Yang, Xuejie Peng, Weiwei Luo, Wen Wang, Ding Xu, Zixiang Huang, Wenlong Sun, Lei Zhao, Qin Yan, Lei Sun, BaoSheng Feng

**Affiliations:** GeneMind Biosciences Co., Ltd., Shenzhen 518000, China

## Abstract

High-throughput sequencing is essential in modern biological research, yet low-complexity sequences remain challenging as they form structurally complex, non-canonical (non-B) DNA conformations that impede sequencing enzyme read-through. This leads to a long-standing trade-off: maximizing coverage introduces false positives (FP), while stringent filtering causes coverage loss and false negatives (FN). To address this, we developed CMS (Cross Mountains and Seas) on GeneMind sequencing platforms by optimizing its chemistry and enzymatic systems to traverse these secondary structures with high fidelity. Benchmarking across whole-genome (WGS) and whole-exome (WES) sequencing demonstrates that CMS addresses the trade-off by simultaneously enhancing both coverage uniformity and accuracy, notably achieving an approximately 100-fold reduction in low-coverage bins for WGS and a 70% reduction in FN insertions/deletions (INDELs) within complex non-B regions. Specifically, a synthetic G-quadruplex (G4) motif sequencing experiment demonstrates that CMS maintains a 1:1 strand ratio, effectively handling G4-induced biases where benchmarked platforms exhibit extensive depletion. These findings establish CMS as a reliable technology for the precise characterization of structural-challenging but functional-essential genome regions.

## Introduction

High-throughput sequencing has established a fundamental role in contemporary genomics research^1–3^. However, a noticeable portion of the genome remains difficult-to-sequence, particularly in regions characterized by low-complexity sequences^4–6^. Despite their simplicity at the sequence level, these regions frequently adopt non-canonical (non-B) DNA structures—such as G-quadruplexes (G4s), Z-DNA and hairpins—rather than the classical B-form double helix. These complex secondary structures create biophysical barriers that hinder the sequencing enzymes^4–6^ . Although challenging to sequencing, these non-B DNA motifs are pervasive (for example, accounting for approximately 13% of the human genome^7^) and particularly enriched in promoters, enhancers, centromeres, and telomeres^8,9^. They playing pivotal roles in gene regulation, chromatin organization, DNA replication, and genome evolution^4,10–15^.

if a sequencing platform aim to maintain high coverage in these difficult-to-sequence regions, a large amount of low-quality data will be retained, and consequently sequencing errors will be introduced, resulting an increase in false positive calls. On the other hand, if stringent filtering is applied to ensure data quality, the sequences in these regions are often lost, resulting in poor coverage and an increase in false negative calls. This trade-off indicates that current sequencing technologies often fail to achieve high accuracy and high coverage simultaneously in these difficult-to-sequence regions^16–20^.

In this study, we introduce CMS, a novel sequencing technology. By systematically optimizing sequencing chemistry and key enzymes, we have enhanced the processivity and fidelity of the sequencing reagent when reading through complex secondary structures. To comprehensively evaluate the performance of CMS, we conducted a systematic comparison of mainstream sequencing platforms using both whole-genome and targeted sequencing approaches. Our results demonstrate that CMS addresses the inherent trade-off between coverage and quality, achieving a simultaneous reduction in both false positive and false negative variant calls, particularly within complex non-B DNA regions. These findings establish CMS as an effective sequencing technology for the comprehensive characterization of genomic structures and their functional roles in biological and pathological processes.

## Results

### Sequencing Data Acquisition and Quality Control

This study constructed six PCR-free DNA duplicate libraries for three standard samples (HG001-HG003, two libraries for each). These libraries were sequenced on three platforms, two repeated sequencing runs on the GeneMind SURFSeq™ 5000 platform with CMS chemistry, one sequencing run on the NovaSeq™ X (NX) platform, and one sequencing run on the DNBSEQ™-T7 (T7) platform. All data were acquired in Pair-End 150 mode, see Methods for detailed procedures for library construction and sequencing. By merging the data from the two libraries originating from the same standard sample to constitute one replicate, we ultimately obtained two replicates per sample for CMS and one replicate per sample for each of the NX and T7 platforms (all derived from the three standard samples). Notably, CMS delivered high quality bases with an average Q40 rate of 97.3%, which is higher than that of the NX (94.8%) platform. Furthermore, 69% of bases from CMS achieved base quality of Q50 (**Supplementary Tables 1**), slightly below the official production specification which is 70% Q50 bases, due to the use of preliminary version of sequencing reagents.

To evaluate performance of CMS in targeted sequencing, WES libraries were constructed for five benchmark samples (HG001–HG005) and sequenced using CMS and the NX platform. The Q40 rate of CMS averaged 98.2%, still higher than that of the NX platform (96.2%); the Q50 rate reached approximately 75.6%, consistently outperforming benchmarked platforms in high-fidelity base calling (**Supplementary Tables 2**).

### Improvement in extremely low coverage regions

To exclude the impact of non-sequencing factors, such as alignment errors or incomplete genome assembly, we first performed the analysis using the high-confidence regions (GRCh37) provided by GIAB, which collectively span 2.31 Gbp. We divided these high-confidence regions into 50-bp bins, calculated the sequencing depth for each bin, and then normalized this depth by dividing it by the median bin depth of each replicate, yielding a normalized depth for inter-replicate comparison.

For alignment and coverage analysis, the sequencing data output was normalized to 2,000M reads per replicate, yielding comparable median depths across all human genome samples, ranging from 90.2x to 101.9x (**Figure 1a**). Accordingly, the thresholds of 0.1x, 0.3x, and 0.5x of the median correspond to absolute sequencing depths of approximately 10x, 30x, and 50x, respectively. To evaluate whole-genome coverage consistency, we analyzed the Coefficient of Variation (CV) of normalized depths. CMS notably exhibited the lowest overall CV among the three platforms; statistical analysis revealed that the uniformity of CMS was higher than that of the NX platform (one-tailed t-test, *P* < 0.01) (**Figure 1b**). These results indicate that CMS provides the highest level of coverage uniformity across the whole genome.

**Figure 1.**
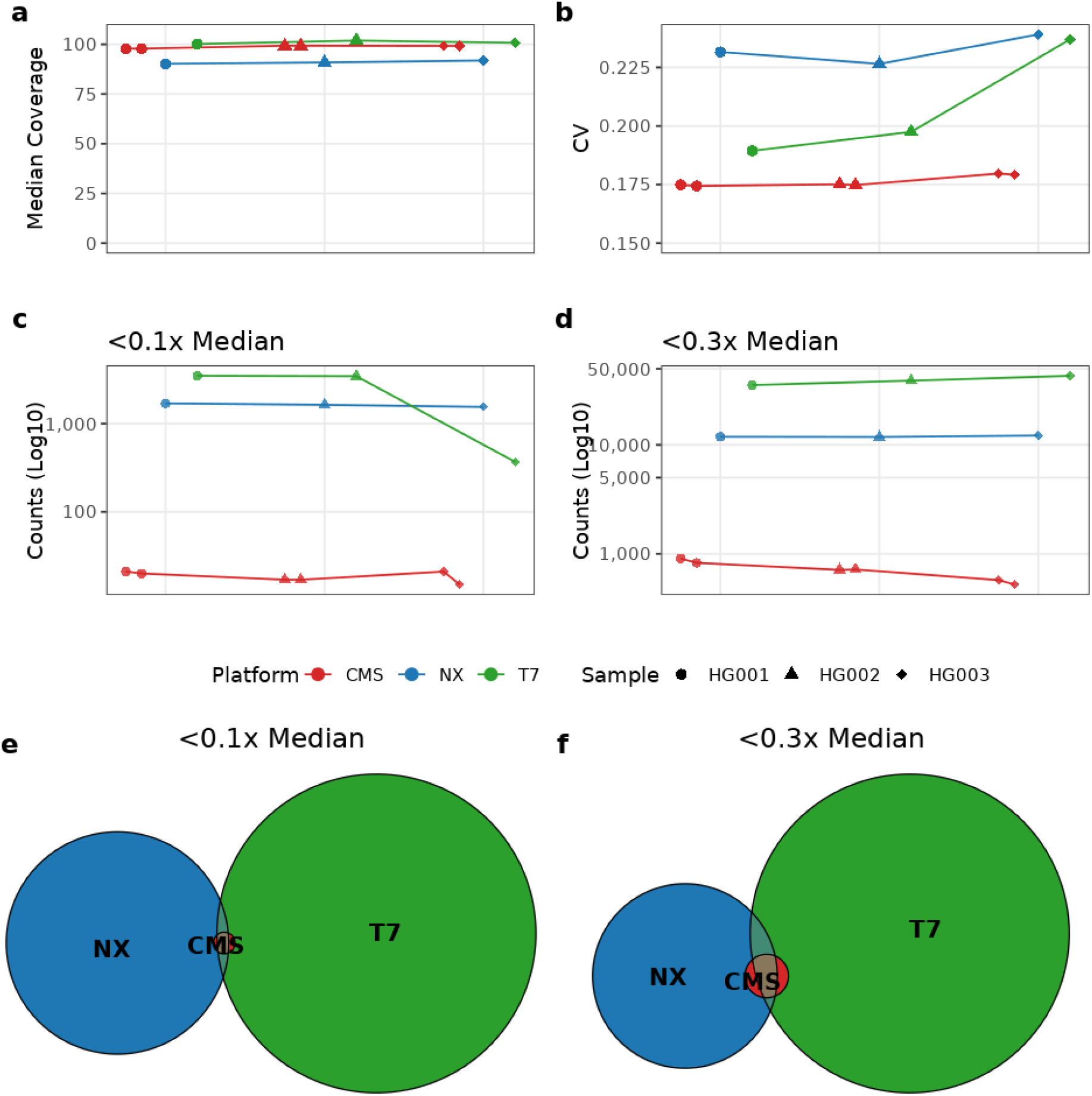
Low-coverage bins across sequencing platforms and samples. Sequencing results for three DNA samples (HG001-HG003) were obtained from three platforms: CMS (two replicates per library), NX (one run), and T7 (one run). The analysis was performed on 50 bp bins within the GIAB high-confidence regions (GRCh37). (a) Median normalized sequencing depth of the 50 bp bins. (b) Coefficient of Variation (CV) of the normalized bin depths, a measure of whole-genome coverage uniformity. (c) Number of low-coverage bins with a normalized depth below the 0.1x median threshold. (d) Number of low-coverage bins with a normalized depth below the 0.3x median threshold. (e, f) Venn diagrams illustrating the overlap of low-coverage bins among the three platforms at the 0.1x (e) and 0.3x (f) normalized depth thresholds.

To better reflect the sequencing performance in difficult-to-sequence regions, which generally have very low depth, we also quantified the number of bins per replicate that had a depth below a defined threshold. As shown in **Figure 1c** and **Supplementary Table 3**, CMS reduced the number of low-coverage (0.1x median) bins to a very low level (averaging 18.5 bins). In contrast, the NX platform (averaging 1623) and the T7 platform (averaging 2209 bins) contained thousands of low-coverage bins. This implies that CMS improved coverage in extremely difficult-to-sequence regions by approximately 100-fold comparing to the NX and T7 platforms. When the threshold was elevated to 0.3X, CMS consistently maintained an extremely low number of low-coverage bins (about 500–900), whereas the benchmarked platforms NX and T7 retained low-coverage bins as high as approximately 12,000 and 40,000, respectively (**Figure 1d** and **Supplementary Table 3**).

To further investigate the platform-specific coverage preferences among different sequencing technologies, we conducted a Venn diagram overlap analysis on the low-coverage bins at the 0.1x and 0.3x thresholds, respectively (**Figure 1e, f** and **Supplementary Table 4**). The results demonstrate that the benchmarked NX and T7 platforms exhibit clear platform-specific low-coverage bins, with a relatively low proportion of shared low-coverage bins. For HG001 at the 0.1x threshold, platform-specific bins accounted for 98.3% (1,665/1,693) and 99.1% (3,451/3,481) of the total for the NX and T7 platforms, respectively; In contrast, the CMS platform showed an extremely few (only 5) unique low-coverage bins. When the threshold was set to 0.3x, this trend was still marked: the unique low-coverage bins for both the NX and T7 platforms exceeded 10,000, while below 250 for CMS. More overlap comparisons for HG002 and HG003 among platforms are provided in **Supplementary Table 4**.

These findings demonstrated that CMS sequencing chemistry can effectively address a wide range of regions with extremely low coverage where benchmarked platforms suffer data loss, which is offering a reliable data foundation for the precise characterization of difficult-to-sequence regions.

### Substantial Coverage enhancement in Non-B regions

To further evaluate the coverage preference of CMS technology across different genomic contexts, we analyzed the normalized depth density distribution for the whole genome (Genome), all non-B DNA structure regions and each specific subtype. As shown in **Figure 2**, CMS exhibits high uniformity in coverage distribution across all examined genomic regions. At whole-genome level, its depth distribution curve is strictly centered around a normalized depth of 1.0, and the curve shape is sharper and more symmetric than those of the NX and T7 platforms (**Figure 2a and 2b**). This is highly consistent with the previous results showing a minimal count of low-coverage bins across the whole genome (**Figure 1c, d**). In contrast, both the NX and T7 platforms show a clear “left tailing” phenomenon in both whole-genome and non-B regions, which is particularly evident for the T7 platform. This means that the low-coverage depth interval (e.g., normalized depth <0.5x median) occupies a larger area, reflecting a widespread depth drop-out in complex regions for other platforms.

**Figure 2.**
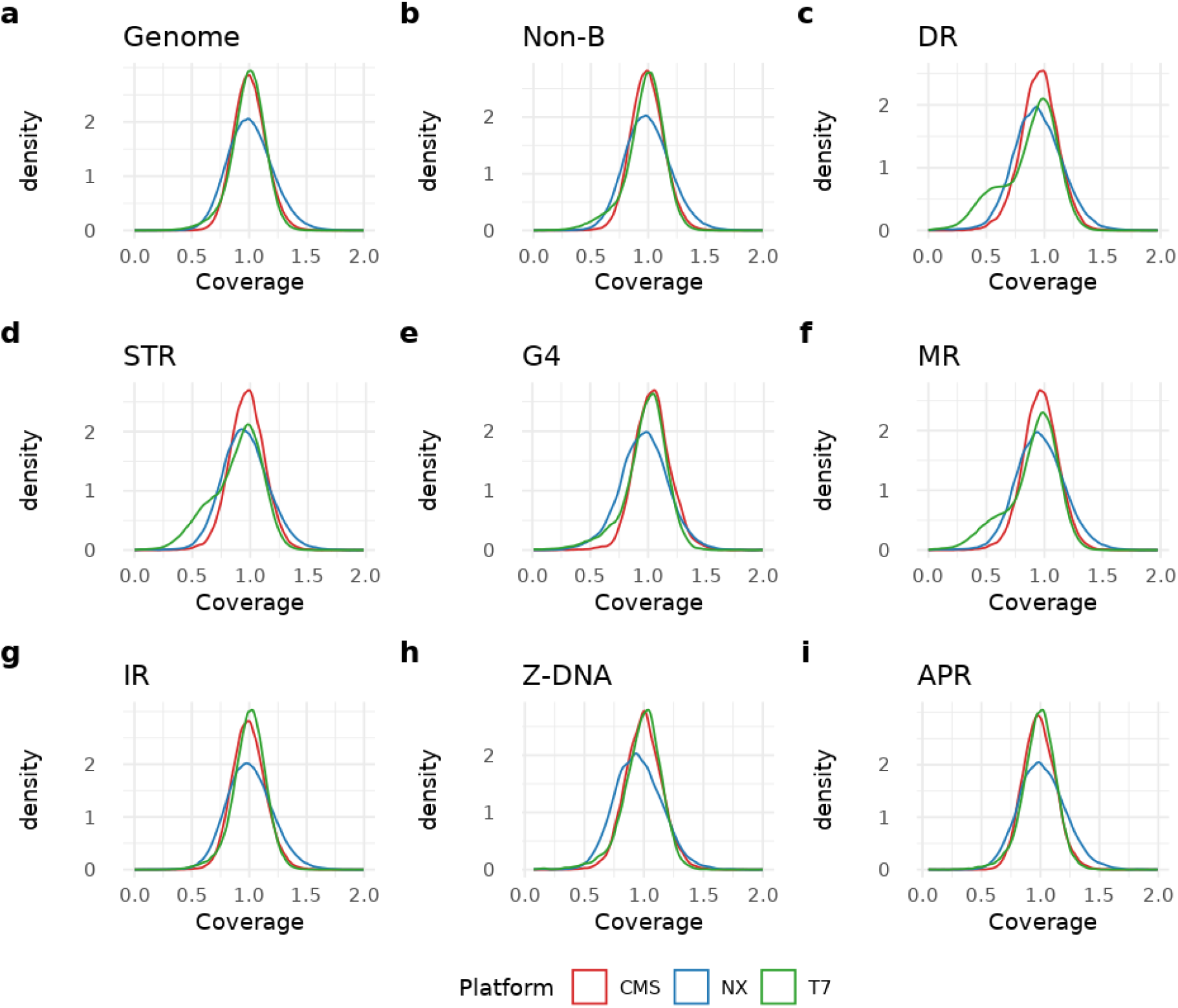
Coverage distribution across genomic and non-B DNA regions. **(a–b)** Normalized coverage density for the whole genome **(a)** and aggregated non-B DNA regions **(b). (c–i)** Distribution profiles for specific motifs: direct repeats **(c)**, short tandem repeats **(d)**, G-quadruplexes **(e)**, mirror repeats **(f)**, inverted repeats **(g)**, Z-DNA **(h)**, and A-phased repeats **(i)**. The CMS platform (red) consistently shows the narrowest peaks centered at 1.0, indicating higher uniformity across complex structures compared to the NX (blue) and T7 (green) platforms.

The advantage of CMS is demonstrated by high uniformity across all specific non-B subtypes (**Figure 2c-2i**). The most noticeable advantage is in G-quadruplexes, Direct Repeats, and Short Tandem Repeats, where the number of low-coverage bins remains at an extremely low level. These findings indicate that the influence of complex non-B secondary structures on the CMS sequencing process is reduced, leading to a substantial improvement in the completeness of genome coverage.

### Performance in Variant Calling Accuracy across Genomic Contexts

To evaluate the sequencing quality, we conducted a comprehensive benchmark of variant calling accuracy for single-nucleotide variants (SNVs) and small INDELs across the whole genome, non-B DNA regions, and low-coverage regions (defined as <0.3x or 0.5x median depth). The systematic evaluation demonstrates that CMS successfully addresses the conventional trade-off, achieving a simultaneous improvement in both genomic coverage and variant-calling fidelity by simultaneously reducing both FP and FN counts (**Figure 3** and **Supplementary Table 5**).

**Figure 3.**
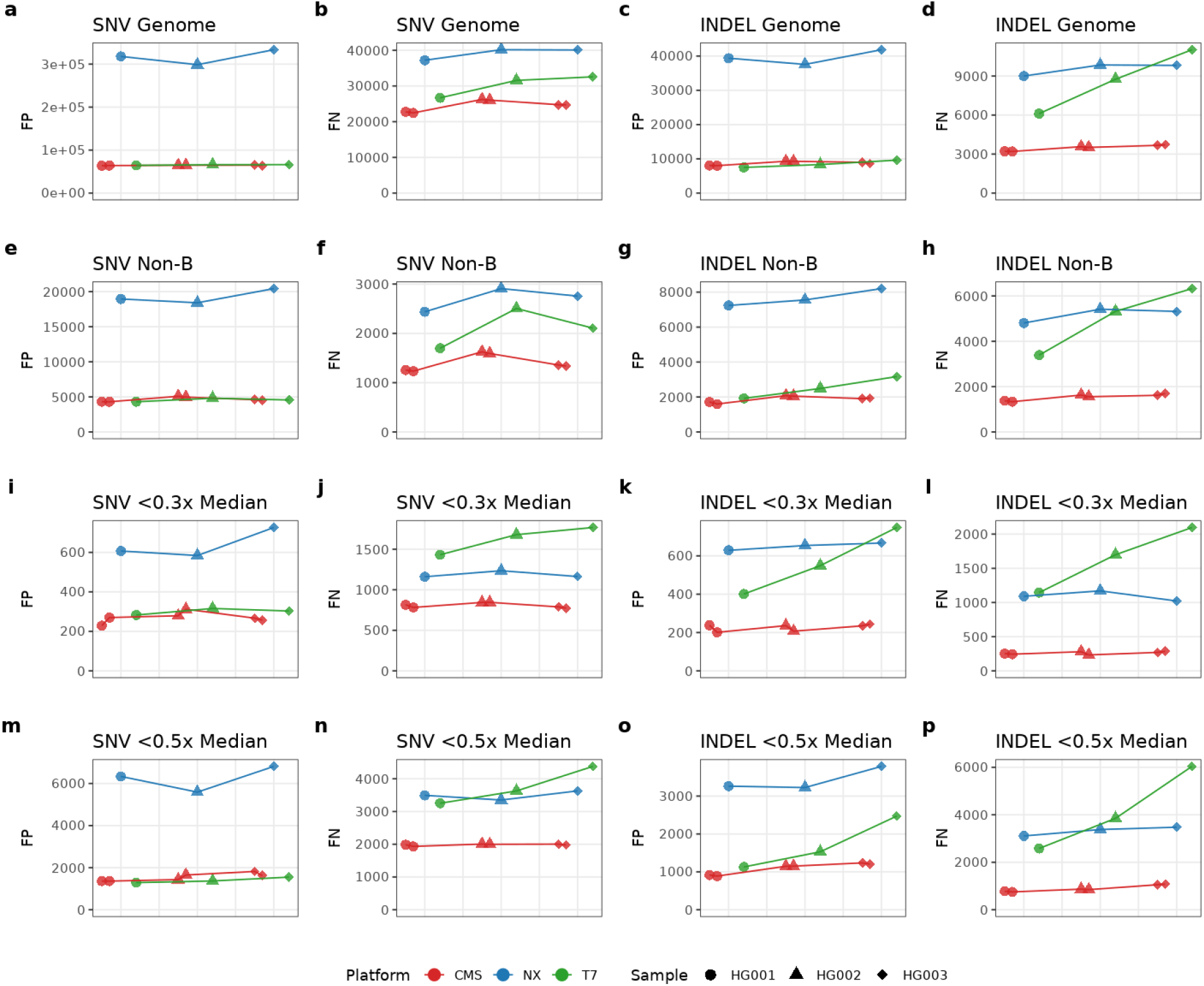
Systematic benchmarking of variant calling accuracy across genomic contexts. (**a-h**) SNV and INDEL performance in whole-genome and non-B DNA regions. Panels (**a, b, e, f**) show False Positive (FP) and False Negative (FN) counts for Single-Nucleotide Variants (SNVs), while panels (**c, d, g, h**) show corresponding metrics for INDELs. CMS consistently achieves lower error rates, particularly in complex non-B DNA motifs. (**i-p**) Accuracy within low-coverage intervals. Performance is evaluated within genomic bins where sequencing depth falls below 0.3x (**i-l**) and 0.5x (**m-p**) of the median—corresponding to approximately 30× and 50× absolute depth, respectively. These intervals comprise the union of low-coverage bins across all platforms to ensure a standardized comparison across identical genomic regions.

In the whole-genome evaluation, the benchmarked platforms confirmed the existence of the trade-off, demonstrating their respective limitations. For SNV detection, CMS achieved the highest recall rate, with an average FN count of 24,493—lower than those of the T7 (30,272) and NX (39,150) platforms. In terms of precision, CMS’s average FP count (64,386) was substantially lower than that of the NX platform (316,645) and slightly lower than that of the T7 platform (65,548). Regarding INDEL performance, CMS’s average FN count (3,480) represented an approximately 60% reduction compared to the T7 (8,631) and NX (9,557) platforms. While the T7 platform’s FP count (8,483) was marginally lower, CMS (8,744) achieved a more than 4.5-fold reduction in false positives relative to the NX platform (39,597).

The performance advantage of CMS was even more pronounced in non-B DNA regions. The average INDEL FN count for CMS (1,400) represented a reduction of approximately 73.0% and 72.1% compared to those of the NX (5,180) and T7 (5,012) platforms, respectively. Similarly, the average INDEL FP count for CMS (1,882) was more than 4-fold lower than that of the NX platform (7,664) and also notably lower than that of the T7 platform (2,526).

When considering the union of all bins with <0.3x median depth across all platforms and replicates (**Figure 3i-l**), the CMS platform’s advantage was more pronounced, leading in all core metrics. In these most challenging regions, CMS demonstrated the strongest performance, achieving the lowest average values for all four key metrics (FN and FP for both SNV and INDEL). For example, its FN INDEL count (261) represented an 84.2% and 76.1% reduction compared to those of the T7 (1,647) and NX (1,094) platforms, respectively, confirming that CMS maintains a very high INDEL recall rate even under low-coverage conditions that challenge other platforms. Consistent performance was observed at the <0.5x median depth threshold as well (**Figure 3m–p**).

### Coverage Evaluation Using T2T Genome and Mitochondrial Genome

While GIAB high-confidence intervals are the gold standard for benchmarking, they often exclude complex non-B DNA regions that CMS has been specifically optimized to address. To minimize the impact of reference genome limitations and mapping artifacts, we used the HG002 T2T (Telomere-to-Telomere) reference assembly to evaluate genome coverage.

The CMS platform demonstrated high genomic coverage and yielded the fewest low-coverage bins. At the <0.1x median coverage threshold (Figure 4a), CMS showed a reduction of approximately 12-fold and 6-fold compared to those of the T7 (49,574) and NX (25,052) platforms, respectively. This performance continued at the <0.3x threshold (Figure 4b), where CMS (140,559 bins) achieved approximately 3.1-fold and 3.8-fold reductions in low-coverage bins compared to the NX (434,387) and T7 (530,037) platforms, respectively (**Supplementary Table 6**).

**Figure 4.**
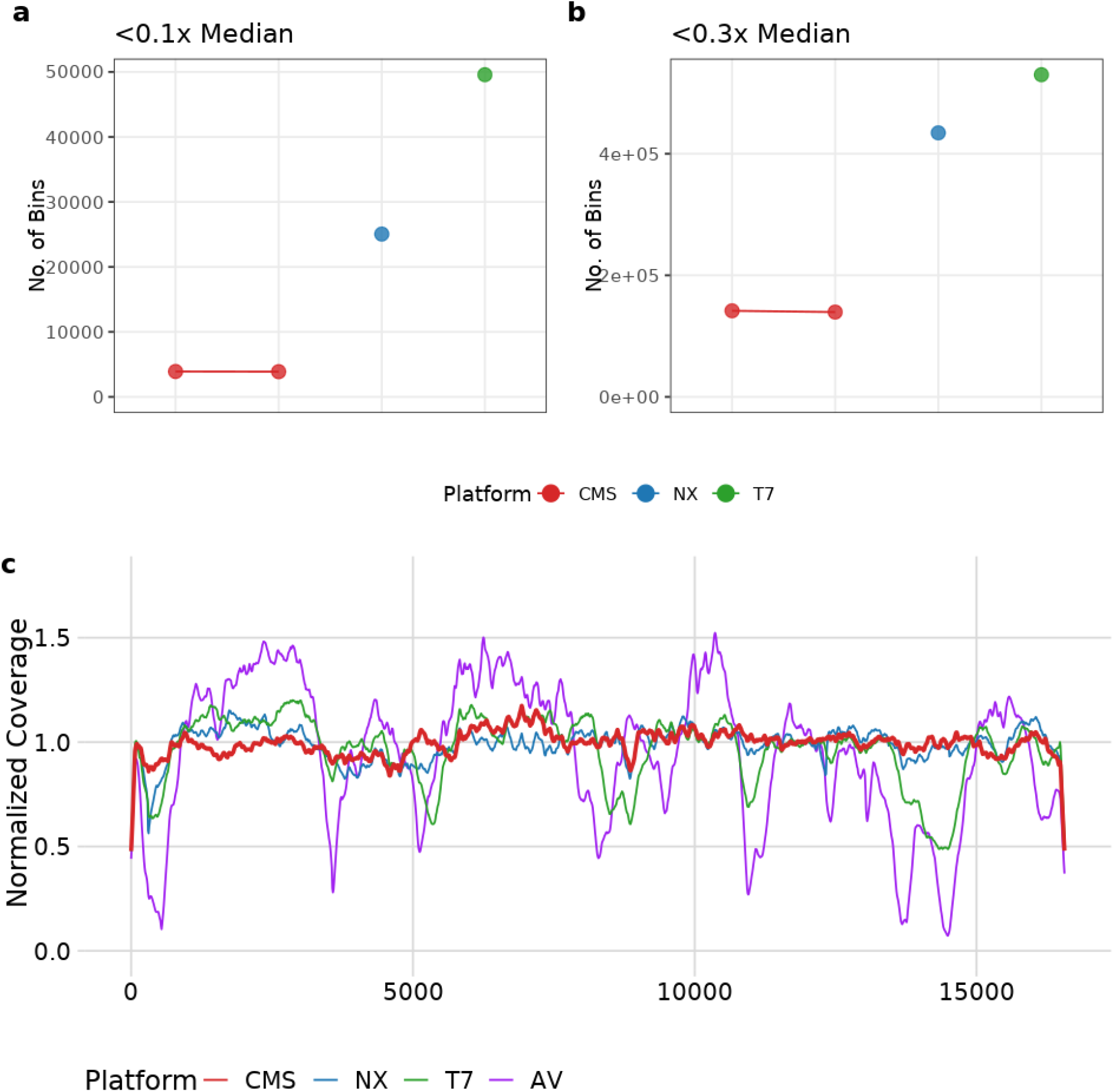
Assessment of Coverage Depth and Uniformity for HG002 T2T and Mitochondrial Genomes. **(a–b)** Number of low-coverage bins (<0.1x and <0.3x of median depth) across the T2T genome for CMS, NX, and T7 platforms. **(c)** Longitudinal normalized coverage across the mitochondrial DNA. Note that AV platform data (purple) is derived from the CHM13 cell line, while all other platforms utilized the HG002 sample. CMS demonstrates the most consistent coverage profile across both nuclear and mitochondrial targets.

Given the extensive use of the mitochondrial genome in studies of population evolution and genetic diseases, we further compared the coverage uniformity of each platform in this region. We divided the mitochondrial genome into 10-bp bins, calculated the sequencing depth for each bin, and normalized the depth by its median for inter-platform comparison. As shown in **Figure 4c**, the CMS platform clearly exhibits the best coverage uniformity on the mitochondrial genome, outperforming both the NX and T7 platforms. We also included sequencing results from the Element AVITI™ (AV) platform on the CHM13 cell line for comparison. This AV dataset showed substantial coverage fluctuation, which may be related to its unique sequencing chemistry^21^.

### Performance Evaluation in Targeted Sequencing

To evaluate the performance of CMS in targeted sequencing, Whole Exome Sequencing (WES) was performed on five benchmark samples (HG001–HG005). Across all replicates, CMS demonstrated improved accuracy compared to the NX platform, achieving a concurrent reduction in both FP and FN rates for SNVs and INDELs (**Figure 5a–d and Supplementary Table 7**). Notably, CMS maintained a consistently lower FN rate for INDELs across all samples, reflecting its robustness in detecting variants within complex exome intervals (**Figure 5d**). The distinction between WGS and targeted sequencing is critical for assessing motif-driven biases. In WGS, random DNA fragmentation creates a “dilution effect,” where only a subset of fragments covering a locus may contain a complete difficult-to-sequence motif at a position that hinders processivity of sequencing reagent. In contrast, targeted sequencing focuses on fixed intervals, resulting a “concentration effect”: if a complex motif is situated within a target region, a higher proportion, in some cases, nearly all of the enriched fragments must be read through. This concentration provides a sensitive window to observe how CMS addresses sequencing obstacles that often lead to coverage deficiencies in other platforms. At a 1.5-fold depth threshold, CMS successfully recovered 2,042 bins that were deficient in the NX platform—approximately 18 times more than the 116 bins where the NX platform showed a depth advantage (**Figure 5e**). Even at a stringent 2.0-fold threshold, CMS resolved 75 coverage-deficient bins compared to 16 for the NX platform (**Figure 5e**). For example, as shown in Figure 5f, CMS achieved smooth, continuous read-through across a G-rich gene region (*KMT2D*), while the NX platform exhibited a sharp decline in coverage.

**Figure 5.**
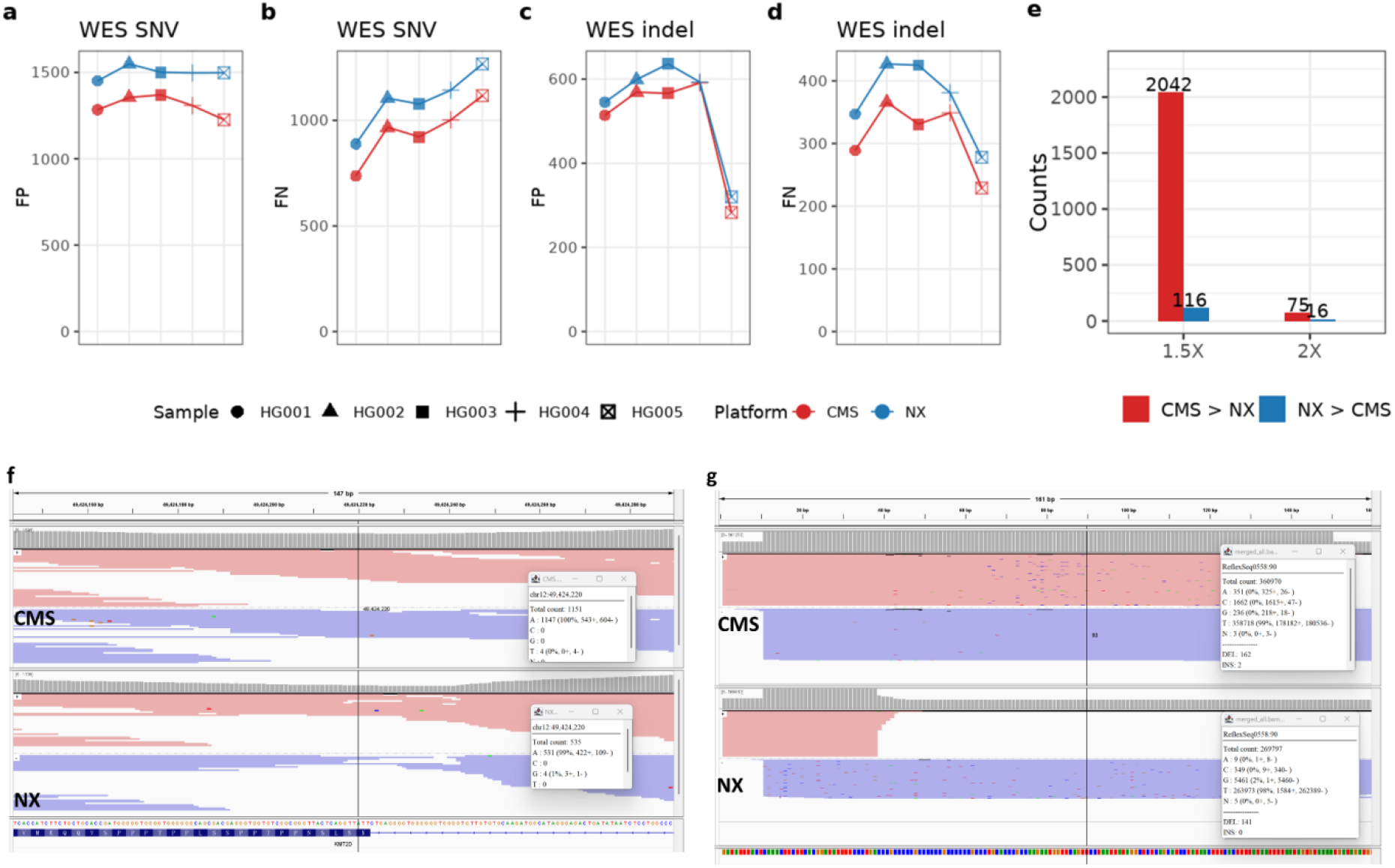
Evaluation of CMS performance in targeted sequencing. (a–d) Comparison of SNV and INDEL calling performance (FP and FN counts) between CMS and NX platforms across five benchmark samples (HG001–HG005) using the WES V6 capture kit. CMS (red) consistently shows lower error rates than the NX platform (blue). (e) Quantification of depth-deficient bins within capture intervals. Bar plots show the count of bins where one platform exhibited a 1.5-fold or 2.0-fold depth advantage over the other. CMS recovered greatly more coverage-deficient bins than the NX platform. (f-g) IGV visualization of a G-rich region in WES and a synthetic sequence. The CMS platform (top) demonstrates smooth and continuous read-through, whereas the NX platform (bottom) shows a sharp decline in coverage.

This capability is further illustrated in a synthetic single-target experiment (**Figure 5g**), where a sequence containing a G4 motif was applied to assess strand-specific performance. The NX platform exhibited extreme strand bias, with a forward-to-reverse strand ratio of approximately 1:166 (1,584 vs. 262,389 reads). This extensive depletion of the forward strand indicates a severe sequencing impediment when the G4 structure serves as the template. In contrast, the CMS platform maintained a balanced 1:1 ratio (178,182 vs. 180,536 reads) with high-quality read-through across the structure. These observations indicate that CMS effectively addresses sequencing stalling induced by complex secondary structures, ensuring unbiased representation in motif-enriched targeted regions.

## Discussion

The core innovation of the CMS platform lies in the systematic optimization of sequencing chemistry and key enzymes, fundamentally enhancing the processivity and fidelity of sequencing reagent when reading through complex non-B DNA structures. This achievement is evident in its coverage performance, which showed an approximately 100-fold reduction in low-coverage bins compared to benchmarked platforms for WGS. Furthermore, the platform exhibited high uniformity across all non-B DNA subtypes (e.g., G4, Z-DNA), validating that CMS’s strategy successfully mitigates structural interference, which is further confirmed by the single-target sequencing experiment.

The second central finding is that CMS successfully addressed the inherent trade-off between high coverage and low false-positive rates. Importantly, this improvement stems from the systematic optimization of the fundamental sequencing chemistry, rather than relying on stringent post-sequencing data filtering. By improving the quality of the raw sequence data, CMS achieves greater variant-calling fidelity by reducing both false positives and false negatives. The advantage in sequencing accuracy is particularly evident where conventional sequencers struggle, such as in non-B regions, where CMS demonstrated stronger ability in detecting INDEL variants. Furthermore, in the extremely low-coverage region (<0.3x median), CMS led in all core variant metrics, confirming a very high INDEL recall rate even in the most challenging genomic contexts.

Despite the substantial improvements achieved by CMS in mitigating sequencing bias in non-B DNA, as a short-read technology, CMS still operates within certain limitations. The short read length (e.g., PE150) inherently restricts the resolution of highly repetitive regions, such as human telomeres and centromeres, where long-read sequencing technologies can achieve reads spanning several megabases. In these complex regions, even perfectly sequenced short reads are subject to considerable alignment challenges, impeding complete assembly and comprehensive variant detection. Furthermore, the current study primarily focuses on the human genome, which may limit the generalizability of these findings to more genetically diverse populations or to organisms with larger, more complex genomes.

Regarding the bioinformatic pipeline, variant calling in this study was performed using NVIDIA Parabricks ^22^, a GPU-accelerated implementation of the GATK HaplotypeCaller ^23^, rather than neural network-based tools such as DeepVariant ^24^. While DeepVariant is capable of demonstrating the advantages of CMS (data not shown), its models are often trained on specific benchmark datasets, which may preclude a fully objective evaluation across the entire HG001–HG005 sample set. In contrast, the non-learning-based HaplotypeCaller algorithm more faithfully reflects the direct impact of improved raw sequencing data quality on variant calling performance, ensuring that the observed gains in accuracy are driven by the chemistry and enzymatic optimizations of the CMS platform itself.

The CMS technology’s capability to characterize complex genomic regions presents several avenues for future development. As a high-performance short-read platform, CMS provides new possibilities for Structural Variation (SV) detection, particularly when integrated with longer read modes (e.g., PE500) to further enhance the utility of short reads in complex genomic analysis. Furthermore, these capabilities extend to plant and animal genomics, where high non-B DNA content often complicates genomic resolution and functional research.

Our preliminary assessments indicate that PCR-based libraries exhibit a higher INDEL error burden compared to PCR-free alternatives, pointing toward systematic biases introduced during PCR amplification. Applying the enzymatic and chemical optimizations of CMS to PCR steps during library construction represents a critical next step. Integrating these advancements across the entire workflow—from library preparation to data acquisition—will provide high-fidelity solutions for clinical diagnostics and oncology, enabling highly precise characterization of the functional genome.

In summary, CMS enhances coverage uniformity and accuracy by mitigating the traditional coverage-quality trade-off. While these results demonstrate its potential for non-B DNA characterization, they are based on a limited scale of library protocols and sequencing runs. Further validation is required to confirm its broad applicability across diverse genomic contexts.

## Materials and Methods

### DNA Samples and Library Construction

Reference genomic DNA samples (HG001-HG003) were utilized for Whole Genome Sequencing (WGS) comparisons. Six PCR-free DNA libraries were constructed for the three standard samples (two per sample). A single library was prepared for each DNA sample (HG001-HG005) using the WES V6 capture kit according to the manufacturer’s instructions.

### Platforms and Sequencing

All samples were sequenced in Pair-End 150 bp (PE150) mode. For WGS evaluation, performance was compared across three platforms: the SURFSeq™ 5000 (utilizing CMS™ chemistry), the NovaSeq™ X (NX), and the DNBSEQ™-T7 (T7). Two replicates were generated for each sample on the CMS platform, with one replicate obtained from each of the NX and T7 platforms. For WES evaluation, the prepared libraries were sequenced on both the CMS and NX platforms to ensure a direct performance comparison.

### External Data Resources

High-confidence benchmark^19^ VCF and BED files (version NISTv4.2.1) for small variants of GRCh37 were obtained from the Genome in a Bottle (GIAB) consortium (https://ftp-trace.ncbi.nlm.nih.gov/ReferenceSamples/giab/release). The HG002 T2T genome assembly^25,26^ was retrieved from the Human Pangenome Resource (https://s3-us-west-2.amazonaws.com/human-pangenomics/T2T/HG002/assemblies/hg002v1.1.mat_Y_EBV_MT.fasta.gz). A corresponding mappability file was utilized to exclude regions with alignment ambiguities, defining the final intervals for coverage comparison. For additional benchmarking, FASTQ data for the CHM13 cell line from the Element Biosciences AVITI™ platform (Cloudbreak UltraQ) was accessed via their resource portal (https://www.elementbiosciences.com/resources/human-whole-genome-sequencing-chm13-cell-line-cloudbreak-ultraq).

### WGS Coverage Evaluation

The GIAB high-confidence regions were partitioned into 50-bp genomic bins. The sequencing depth for each bin was calculated and normalized by the median bin depth of the corresponding replicate (Normalized Depth = Bin Depth / Median Bin Depth). Coverage uniformity was quantified using the Coefficient of Variation (CV) of the normalized depths. To specifically assess regions of coverage loss, we quantified the frequency of low-coverage bins. These were defined using multiple thresholds, for example, a threshold of 0.1x median identifies bins where the normalized depth falls below 0.1, which represents a substantial depletion of reads. Furthermore, coverage distributions were stratified and analyzed across various non-canonical (non-B) DNA structures to evaluate the platform’s capability to resolve structurally complex genomic regions. Non-B motifs^10^ of the GRCh37 genome were downloaded from https://nonb-abcc.ncifcrf.gov/, including Direct Repeats, Short Tandem Repeats, G-quadruplexes, Mirror Repeats, Inverted Repeats, Z-DNA and A-phased repeats.

### WGS Variant Calling Evaluation

To account for the variations in effective sequencing depth caused by disparate duplication rates across the benchmarked platforms—most notably the higher duplicate read output of the NX platform—a standardized depth normalization protocol was employed. For each replicate, 2,000 million raw reads were aligned to the reference genome, and duplicate reads were identified and marked. The median sequencing depth was subsequently calculated using only the unique (non-duplicate) reads. To facilitate a rigorous and equitable comparison of variant calling accuracy, the data were proportionally downsampled from the unique read pool to achieve a standardized target mean depth of 30x for all subsequent SNV and INDEL evaluations.

Variant calling was performed using the NVIDIA Clara Parabricks pipeline, utilizing the GATK HaplotypeCaller algorithm, to generate VCF files. Variant accuracy for SNVs and small INDELs was assessed using RTG Tools vcfeval^27^ against the GIAB high-confidence truth set. Performance metrics, including False Positives (FP), False Negatives (FN), Recall, and Precision, were calculated within the high-confidence regions.

### WES Coverage Evaluation

To evaluate coverage performance in WES, HG001–HG005 reads were downsampled to 140M per sample and pooled to a total of 700M per platform, achieving an average sequencing depth of approximately 2,000x across target regions. Coverage uniformity within the WES V6 capture intervals was assessed using a sliding window approach (20-bp window size, 10-bp step). Per-bin depth was computed using mosdepth (v0.3.2), and normalized depth was subsequently determined by dividing the depth of each bin by the median depth across all bins.

A “depth deficiency” criterion was established to identify regions of pronounced coverage loss. A window was considered deficient on one platform if the normalized depth ratio between the two platforms exceeded a specific threshold (1.5x or 2.0x). To ensure the reliability of this identification, we required the platform with higher depth to maintain a normalized depth > 0.3, while the platform with lower depth exhibited a normalized depth < 0.5. This analysis allowed for the precise identification of coverage-deficient bins and the quantification of each platform’s ability to recover challenging capture targets.

### WES Variant Calling Evaluation

To ensure a standardized and equitable comparison of targeted sequencing performance, data output of each replicate was downsampled to 12 Gbp, corresponding to an average sequencing depth of approximately 100x. These standardized datasets were subjected to alignment and variant calling procedures identical to the methodologies previously described for the WGS evaluations. Variant calling performance was then assessed within the intersection of the WES V6 target intervals and the GIAB high-confidence regions to ensure that the analysis was restricted to high-quality genomic regions covered by the capture kit.

### Synthetic G4 Experiment

A synthetic sequence containing a G4 motif was utilized for a single-target experiment to evaluate read-through capability. The sequence is as follows: GATATTTACTCAAACTTCTTCACGGATATTTTCCCTGACCCGCCCCCGCCCCCCGCCCCCCGCCCTGCACCCGCCAAGAATGGTTCAGTTCCTAATTATAGGCTCTCAGCATCCTATAACTTTCACTTTTATAATTTTAATAGTGTAAAATGATAGATGT.

According to the sequence design, the G4 motif is located on the reverse strand. During sequencing, when the reverse strand serves as the template for read synthesis, the complex secondary structure of the G4 motif hinders processivity of sequencing reagent. This physical impediment results in a substantial reduction of reads mapping to the forward strand, leading to extreme strand bias. Strand-specific performance was evaluated by calculating the forward-to-reverse read ratio across the G4 motif.

## Acknowledgments

This work was supported by the National Science and Technology Major Project of China (Grant No. 2023ZD0505901).

